# Integrating Bottleneck Size into Selection Tests for Biological Diversity Data

**DOI:** 10.64898/2026.07.07.737025

**Authors:** Thi Minh Thao Le, Erida Gjini

## Abstract

Population bottlenecks profoundly shape genetic diversity, but distinguishing stochastic drift from selective pressure requires precise estimation and accounting for bottleneck size. While deep-sequencing data enable inference via frameworks like beta-binomial modeling, integrating these estimates directly into selection tests remains a critical challenge. In this study, based on existing computational approaches, we propose a new method that explicitly incorporates bottleneck size estimates into neutrality tests for biological diversity data. Designed for variant frequency data, our framework accounts for sequencing errors and sampling biases to improve the precision and interpretability of selection signature detection. We validate this framework using previously published *Streptococcus pneumoniae in vivo* experimental data, successfully replicating established fitness results, while uncovering novel genes relevant to infection and pathogenesis. This integrated new model with explicit bottleneck effects narrows down the set of candidate genes under selection and provides a robust, generalizable tool for disentangling drift from selection across a wide range of biological systems.

## 1. Introduction

In infectious disease dynamics and microbial evolution, identifying which pathogenic genes drive host adaptation and virulence is critical for developing targeted therapies and vaccines. However, during host-to-host transmission or colonization of new anatomical niches, pathogen populations frequently undergo severe constrictions, known as transmission bottlenecks (Figure 1). These bottleneck events induce massive genetic drift, causing stochastic shifts in variant frequencies that can easily be misidentified as directional selection by standard analytical tools. Distinguishing genuine selective pressure from stochastic drift therefore remains a fundamental challenge in interpreting high-throughput genomic and experimental diversity data. In classical population genetics, Krimbas and Tsakas (1971) pioneered the use of the variance of allele frequency shifts across multiple independent polymorphic loci to statistically distinguish genetic drift from selective pressure. Bottleneck size estimation methods infer the number of founding individuals that successfully navigate a restrictive event between populations or time points.

**Figure 1:**
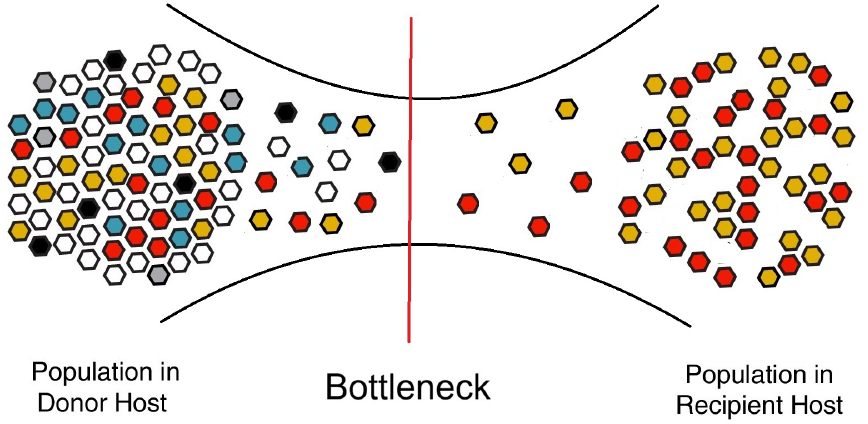
Schematic illustration of a transmission bottleneck. A diverse pathogen population in the donor host passes through a narrow transmission stage, during which only a small subset of individuals successfully founds infection in the recipient host. As a result, the recipient population is initiated by an effective founding population that is much smaller and less diverse than the donor population, so that some variants are lost purely by stochastic sampling while others become overrepresented by chance.

The earliest explicit formalization of population bottleneck sizes and their quantitative effects on heterozygosity and allelic diversity was established by Nei et al. (1975). Prior foundational work (e.g., Wright (1931)) described integral mechanics like genetic drift and effective population size, but did not formalize bottleneck size estimation. By modeling these dynamics, Nei et al. (1975) laid the theoretical groundwork for subsequent estimation methods in evolutionary and conservation genetics. Modern frameworks build upon a consistent extension of this concept; conceptually, both Abel et al. (2015) and Sobel Leonard et al. (2017) define bottleneck size as the effective number of pathogens surviving a population constriction to colonize a new host or anatomical compartment.

While early population genetics literature implicitly applied binomial frameworks (Appendix S1.1) to allele frequency shifts, Sobel Leonard et al. (2017) provided a seminal explicit application in genomic analysis using binomial sampling and beta-binomial extensions. Their likelihood framework models site-specific variant transmission via binomial processes, combining data across sites to estimate global bottleneck size.

While various computational methods exist to estimate the physical size of these bottlenecks, ranging from frequency-based beta-binomial models to coalescent frameworks, integrating these estimates directly into downstream selection tests is rarely done. Most established differential abundance frameworks (such as DESeq2 or MAGeCK) were designed for transcriptomic or large population *in vitro* screens. Because they lack the physical transmission parameters required to bound stochastic variance, applying these methods to *in vivo* bottlenecked data frequently causes them to misinterpret founder-effect drift as biologically significant selection.

To address this limitation, we build upon existing modeling frameworks for inferring bottleneck sizes and propose a new, integrated Bottleneck-Neutrality test. Our method explicitly incorporates Dirichlet-multinomial bottleneck size estimation directly into a likelihood-ratio neutrality test. By establishing a mathematically bounded null hypothesis of genetic drift based on physical transmission constraints, our approach structurally isolates true genotypic selection from stochastic loss.

To illustrate the applicability of our method, we validate the BN test using *in vivo* data from the *Streptococcus pneumoniae* CRISPRi-seq study by Liu et al. (2021) as a foundational reference. We demonstrate how this integrated framework improves the precision of selection signature detection compared to traditional regression-based methods, successfully replicating established fitness results while eliminating compositional false positives driven by narrow founder populations.

## 2. Method

### 2.1 Generalizationofbeta-binomialto Dirichlet-multinomial sampling method

We start by recalling that in Sobel Leonard et al. (2017), the size of bottleneck is calculated from the likelihood functions based on the data of variant frequencies (See Table 1):

**Table 1:**
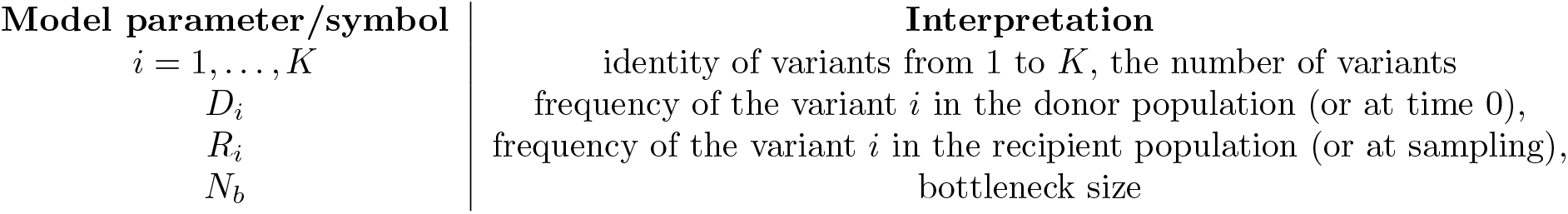
Key parameters and notations in our modeling framework. We assume no selection during transmission (neutrality) and no mutation during the bottleneck.

- For site *i* where the variant is called present:

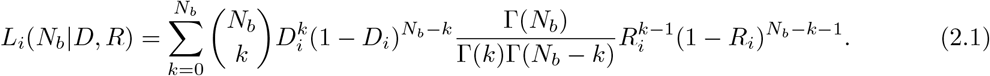
- If the variant *i* is not called under variant-calling threshold: *T* ∈ (0, 1) (frequency below threshold *T*):

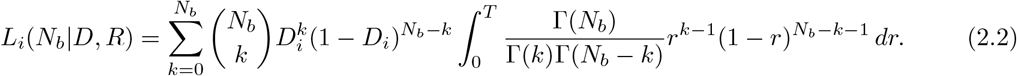

We note that, for the boundary cases *k* = 0 and *k* = *N*_*b*_, the Beta density is technically undefined due to the Gamma function poles at zero. Following Sobel Leonard et al. (2017), these terms are handled via point masses (Dirac delta functions) at *R*_*i*_ = 0 and *R*_*i*_ = 1, respectively. See the proof of (2.1) and 2.2 in Section S1.2.

By treating each site as an independent observation, the total likelihood for a donor–recipient transmission pair is calculated as the product of individual site-specific likelihoods 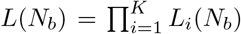Although So-bel Leonard et al. (2017) explicitly acknowledge that this assumption is violated by extensive genetic linkage within influenza virus gene segments, they employ a “data-thinning” strategy to mitigate bias.

Under the assumption of site independence, the beta-binomial sampling method cannot be used to estimate the transmission bottleneck size from compositional data, that is, data in which the frequencies across all sites sum to one. Hence, we extend the beta-binomial model to a multivariate context. For this purpose, we utilize the Dirichlet Multinomial (DM) sampling method to estimate the transmission bottleneck size.

This biological exclusivity is naturally captured by the multinomial component of the model, which replaces the binomial distribution to describe the simultaneous sampling of *K* distinct variants from the donor population. Furthermore, the Dirichlet distribution generalizes the beta distribution by modeling the joint frequency fluctuations (stochastic drift) of all variants on a (*K*−1)-dimensional simplex, ensuring that the sum of variant frequencies remains constrained to unity.

Integrating these components results in the following joint likelihood expression

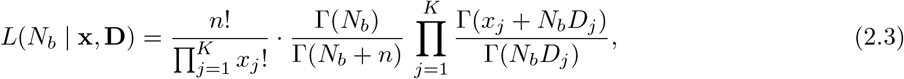

where *n* is the total read count and *x*_*j*_ is the observed count for variant *j* at sampling time (thus 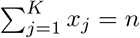 This formulation effectively treats the bottleneck size *N*_*b*_ as the precision parameter of the Dirichlet prior, such that smaller *N*_*b*_ values correctly capture the increased overdispersion and variance associated with narrow transmission bottlenecks.

### 2.2 Integrating bottleneck into selection detection: BN test, a new method

We propose a bottleneck-neutrality test to identify variants under selection, based on the selective neutrality assumption of typical bottleneck models. Unlike classical approaches Liu et al. (2021), that use bottleneck as a quality filter, our framework explicitly incorporates the estimated bottleneck size into selection inference to account for stochastic drift and sampling variance. This test determines whether an individual variant frequency deviates from neutral expectations. Under neutrality, the recipient frequency should fall within the probabilistic range defined by *N*_*b*_ and the initial inoculum frequency; variations exceeding this threshold indicate selective pressure.

#### Data

The data required are read counts of all variants in the inoculum, and read counts of all variants in each host (which is a realization of a stochastic process) after in-vivo colonization at one time point *t*. We treat each host as an independent replicate of the colonization process. Let *D*_*j*_ denote the donor (or inoculum) frequency of variant *j*, with Σ_*j*_ *D*_*j*_ = 1 as before. Let *x*_*m,j*_ be the observed read count of variant *j* in host *m* at time *t* post-colonization, and 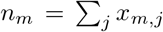 the total read count in that host. Let *R*_*m,j*_ denote the recipient frequency of variant *j* in host *m* at time *t*, then 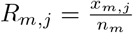 We then proceed in the following steps.

#### 1. Estimate bottleneck size per host

For each host *m*, we fit a Dirichlet Multinomial model to the entire variants’ count vector to obtain an estimated bottleneck size *N*_*b,m*_.

#### 2. Marginal model for a single variant under DM

For each host, under the DM model with concentration *N*_*b,m*_ and mean composition **D**_*m*_, the marginal distribution of *x*_*m,j*_ is Beta–Binomial:

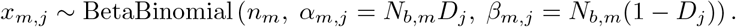

#### 3. Neutrality hypothesis and likelihood across hosts

Under neutrality for variant *j*, the expected recipient frequency *R*_*j*_ equals the donor frequency *D*_*j*_. We write this as *q*_*j*_ = *D*_*j*_. To test neutrality while combining all hosts, we use the joint likelihood across hosts (independence assumption):

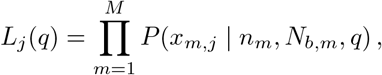

where *P* (*x*_*m,j*_ *n*_*m*_, *N*_*b,m*_, *q*) is the Beta–Binomial probability mass function (pmf). Equivalently, we work with the log-likelihood

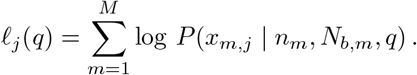

#### 4. Likelihood-ratio neutrality test (LRNT)

For each variant *j*, we compare *H*_0_: *q* = *D*_*j*_ (neutrality, i.e. the frequency in the recipient does not differ much from the donor), and *H*_1_: *q D*_*j*_. Let 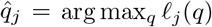. The test statistic is 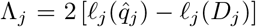, which we compare to a *χ*^2^-distribution to obtain a *p*-value, according to Wilks’ theorem. The proof is in Box 1.

#### 1. Multiple-testing correction

Because many variants are tested, we control the false discovery rate (FDR) across variants using the Benjamini–Hochberg procedure.

##### BOX 1. Proof of Likelihood-Ratio Neutrality Test.

1. For each variant, significance was assessed using a likelihood-ratio test comparing a null model in which the recipient frequency equals the donor frequency (*H*_0_ : *q* = *p*_0_) against an alternative model in which the recipient frequency is estimated freely (*H*_1_ : *q* ≠ *p*_0_).
2. The test statistic is

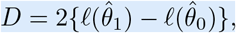

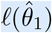 and 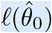 are the maximized log-likelihoods under the alternative and null models, respectively.
3. Under standard regularity conditions, Wilks’ theorem states that, under *H*_0_, the likelihood-ratio statistic converges in distribution to a chi-square random variable with degrees of freedom equal to the difference in the number of free parameters between the two models, that is,

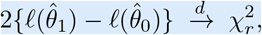

where *r* = dim(Θ_1_) − dim(Θ_0_), where Θ_0_ and Θ_1_ denote the parameter spaces of the two models:
  - Θ_0_: all parameter values allowed under the null model,
  - Θ_1_: all parameter values allowed under the alternative model.

The test compares a full model allowing independent donor and recipient frequencies against a null model where they are constrained to be equal. Since this null hypothesis imposes an equality constraint, the likelihood-ratio statistic asymptotically follows a 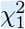 distribution (one degree of freedom).

## 3 Results

### 3.1 Application of BN method to published data in Liu et al. (2021)

Liu et al. (2021) utilized high-throughput CRISPRi-seq to map the *S. pneumoniae* fitness landscape, employing a statistical pipeline that integrated bottleneck quantification with differential expression analysis to isolate genotypic selection from stochastic loss. To differentiate these dynamics, the experimental group (With-Dox) was compared to a parallel, uninduced control group (No-Dox). This CRISPRi-OFF baseline successfully accounted for host-imposed confounding factors, including transmission bottlenecks, immune clearance, and resource competition.

#### Bottleneck quantification originally as a quality filter only

Before assessing gene essentiality, the foundation bottleneck sizes (*N*_*b*_) at 24 and 48 hpi (Liu et al., 2021, Figures 4B and 4C) were calculated using the framework of Abel et al. (2015) (Section S1.3, formula (S2)) based on raw sequencing data from (Liu et al., 2021, Table S7) ^1^. This pronounced constriction indicates that sgRNA loss at 48 hpi was driven primarily by stochastic sampling rather than gene-specific fitness effects. Consequently, the selection analysis was restricted to the 24 hpi, as the extensive loss of library diversity rendered the 48 hpi dataset unreliable for essentiality testing. The replication and recomputation of bottleneck sizes from Liu et al. (2021) is detailed in Supplementary Section S2.2.

#### Statistical Identification of Essential Genes

The core analysis used the DESeq2 package, which fits a generalized linear model (GLM) based on a negative binomial distribution to the sgRNA counts. The primary fitness metric was the log_2_ fold change (log2FC):

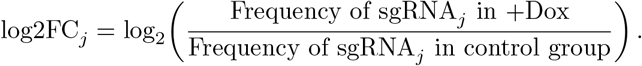

The criteria for essentiality of a sgRNA is log2FC ≤ − 1 and a Benjamini–Hochberg (BH) adjusted *p*-value *p*_adj_ *<* 0.05 ^2^.

#### Differential Fitness and Virulence Factors

To isolate genes specifically required for infection (virulence factors) versus those required for general metabolism, the authors computed the Differential Fitness^3^:

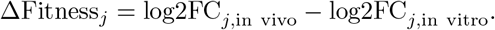

In the following, we apply our BN method to this dataset. We test the neutrality assumption for the observation data at 24 hpi in Liu et al. (2021). For each sgRNA *j*, the donor frequency *D*_*j*_ was estimated from the inoculum (Pre1-for mice in control group and Pre2-for mice in doxycycline-induced group). Recipient data consist of raw counts *x*_*m,j*_ and total reads *n*_*m*_ from each no-dox mouse *m* at 24 hpi.

#### 3.1.1 Result of BN test in mice without doxycycline

For the bottleneck sizes computed for this dataset for use in our Bottleneck-Neutrality test, see Section S2.2. In this control condition, all sgRNAs are expected to behave as neutral. We can still apply the test for the sake of illustration and validity. We find that the test detects 36 sgRNAs which may violate the neutrality assumption, listed in Table 2.

**Table 2:** Identification of 36 sgRNAs deviating from neutrality in no-dox mice at 24 hpi. This table lists sgRNAs that reject the neutrality hypothesis after FDR control (*p*_value_≤0.05). Columns report sgRNAs corresponding the direction of change (up/down in recipient relative to donor).

| Up in frequency ↑ (20 sgRNAs) | Down in frequency ↓ (16 sgRNAs) |
| --- | --- |
| sgRNA1347, sgRNA0406, sgRNA0575, sgRNA1457, sgRNA1296, sgRNA1496, sgRNA954, sgRNA0534, sgRNA0992, sgRNA0814, sgRNA0173, sgRNA0224, sgRNA1166, sgRNA0521, sgRNA1125, sgRNA0204, sgRNA1357, sgRNA0349, sgRNA1249, sgRNA1390 | sgRNA0396, sgRNA0501, sgRNA0492, sgRNA0081, sgRNA0525, sgRNA0461, sgRNA0397, sgRNA0565, sgRNA0302, sgRNA0443, sgRNA0809, sgRNA0808, sgRNA0512, sgRNA0464, sgRNA0272, sgRNA310 |

Under the uninduced (No-Dox) condition, dCas9 is absent and target gene repression cannot occur, meaning each sgRNA functions strictly as a neutral barcode to measure drift and bottleneck effects. Consequently, identifying non-neutral sgRNAs in this setting represents a statistical false-positive effect rather than true biological selection. Given approximately 1,500 tests and a significance threshold of *p*_adj_ = 0.05, about 75 false positives are expected by chance alone (1, 500×0.05 = 75). The 36 detected sgRNAs fall well within this expected range of random fluctuation and do not constitute evidence of systematic biological selection.

If we choose the threshold of rejection for *p*_adj_ 0.001, there still be 9 sgRNAs non-neutral ^4^. Beyond statistical false positives, several biological and modeling factors may explain the apparent non-neutrality of these nine sgRNAs under uninduced conditions: i) Insertion locus effects: Genomic integration near the target gene may disrupt local regulatory elements or chromosomal architecture, inducing fitness costs. ii) Off-target effects: Constitutive sgRNA expression combined with trace dCas9 expression from promoter leakiness can drive unintended repression at partially homologous genomic loci; and iii) Unmodeled bottleneck dynamics: The assumption of a simple, homogeneous transmission event omits the confounding variance introduced by multi-generational stochastic drift, compounding founding population effects, and sequencing noise.

### 3.1.2 Result of BN test in mice with doxycycline

Under dCas9 induction (CRISPRi-ON), target-specific gene repression introduces systematic selection, shifting recipient frequencies away from donor baselines (*q*_*i*_ ≠*D*_*i*_). This selective pressure, compounded by non-target-specific confounding factors—such as CRISPRi metabolic burden, polar effects, and off-target binding—drives widespread deviations from neutrality. Applying our method to this induced context identified a substantially larger pool of sgRNAs violating the null hypothesis, resolving **265 down-regulated** and **966 up-regulated** sgRNAs in the recipient population. The result of non-neutral sgRNAs in the experiment of mice induced by doxycycline are illustrated in the following Figure 2.

**Figure 2:**
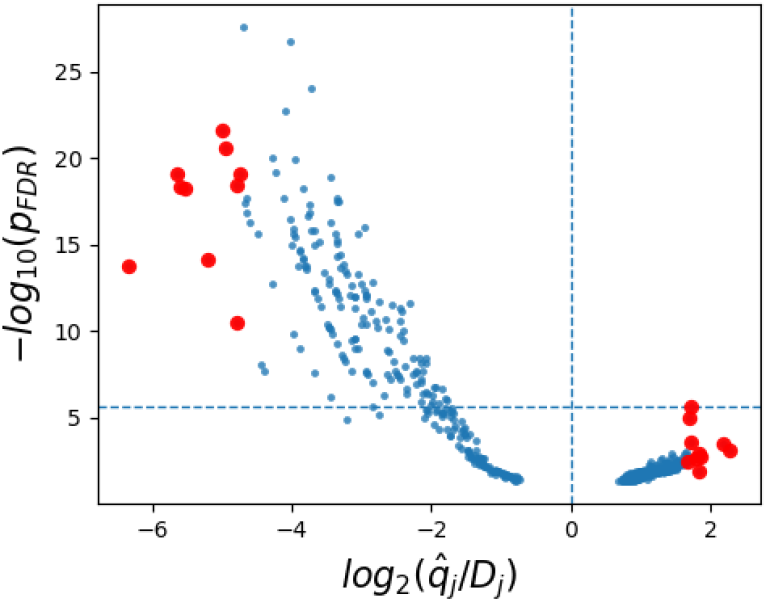
Result of bottleneck-neutrality test applied to sgRNAs in the experimental group in Liu et al. (2021). Each point represents one sgRNA detected to be non-neutral under *p*-value = 0.05. The x-axis shows the log_2_ fold-change-like quantity log 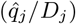 so values *>* 0 and *<* 0 indicate enrichment and depletion relative to donor, respectively. The y-axis shows statistical significance *−* log_10_(*p*_FDR_), where larger *y* corresponds to smaller FDR-adjusted *p*-values (stronger evidence against the null). The vertical dashed line at *x* = 0 marks the no-change boundary, separating negative (“down”/depleted) from positive (“up”/enriched) directions. The ten large red dots on each side of the figure represent the sgRNAs that deviate most strongly from neutrality according to the ratio 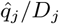, namely those with the smallest and largest values of this ratio. Specifically, the ten data red points on the left are further analyzed in Table 3, and the identities of the ten highest points on each side are presented in Table 4.

In Figure 2, there are two quantities to consider when assessing the neutrality of a variant: the *p*_FDR_ and the ratio 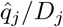, although from this figure we can see that the two are strongly correlated. This means that it is very likely that for an sgRNA that deviates more strongly in magnitude from the 0 border, such deviation is also more likely to be more significant. This means that we can select genes of interest based on either criterion: the ratio and the p-value.

**Table 3:**
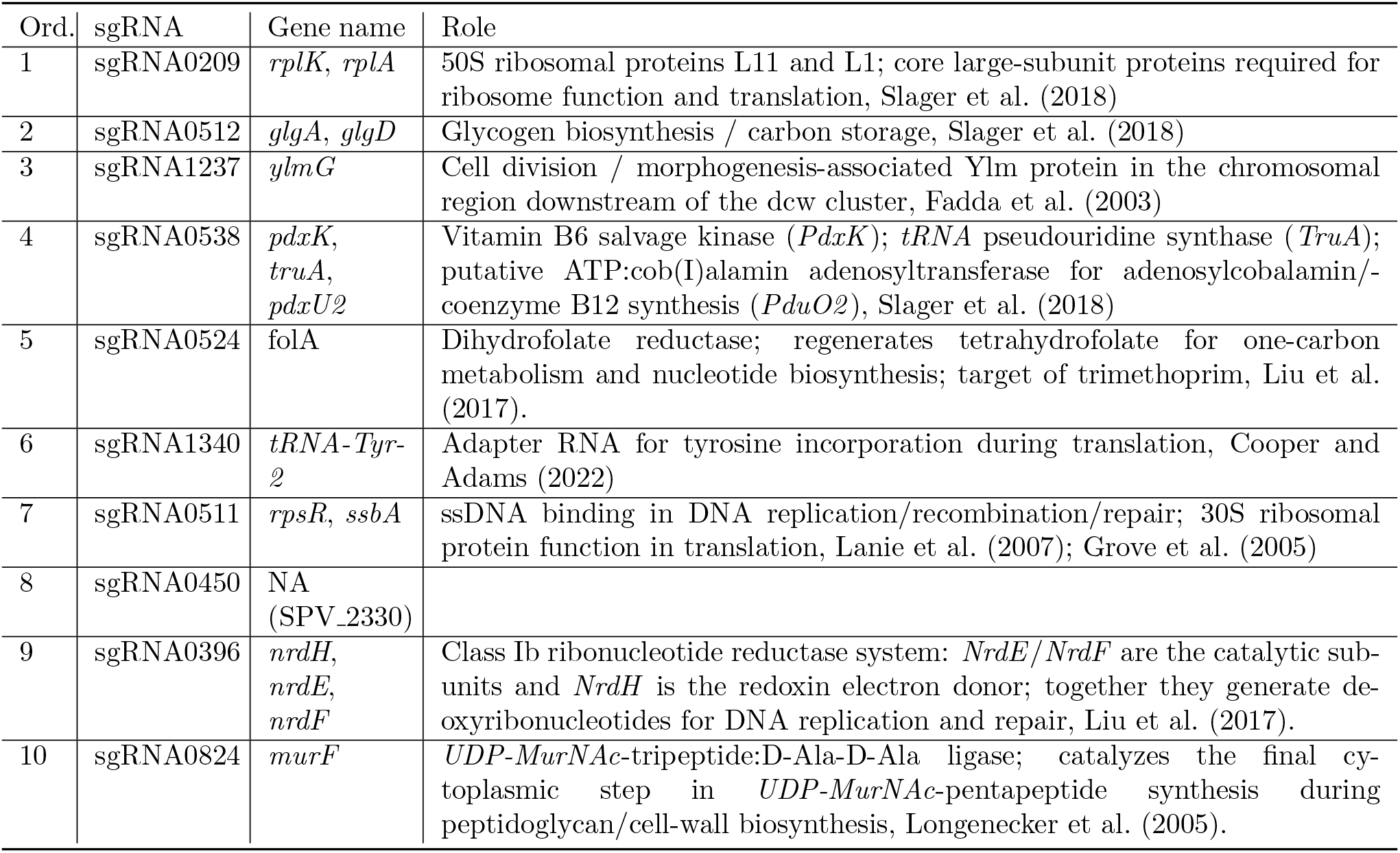
sgRNAs with the smallest ratio 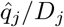 in the bottleneck-neutrality test. (the ten red dots on the left-side of Figure 2) For each sgRNA, the corresponding locus tag, annotated gene, and reported functional role are shown. These sgRNAs represent the strongest depletion from the bottleneck-aware neutral null model, implying strongest signal for selection.

**Table 4:** Some sgRNAs most strongly rejected from neutrality in the with-doxycycline mice at 24 hpi, as determined by the smallest FDR-adjusted *p*-values. This table lists sgRNAs that reject the neutrality hypothesis after FDR control (*p*_FDR_≤0.05) (ten highest points on each side in Figure 2). Columns report sgRNAs corresponding the direction of change (up/down in recipient relative to donor).

| Up in frequency ↑ (10 sgRNAs) | Down in frequency ↓ (10 sgRNAs) |
| --- | --- |
| sgRNA0223, sgRNA0747, sgRNA0212, sgRNA1252, sgRNA1432, sgRNA0483, sgRNA1460, sgRNA1366, sgRNA0406, sgRNA0548 | sgRNA0501, sgRNA0765, sgRNA0860, sgRNA0676, sgRNA1340, sgRNA0511, sgRNA1342, sgRNA0650, sgRNA0302, sgRNA0512, sgRNA0824, sgRNA0513 |

When applying the BN test to the 24 hpi +Dox dataset, the neutrality test identifies a large fraction of guides as deviating from the inoculum, with a pronounced skew toward apparent increases in recipient frequency (966 up-direction vs. 265 down-direction). This imbalance is expected in pooled, sequencing-based screens because the data are compositional: when many sgRNAs drop out, the remaining sgRNAs necessarily occupy a larger share of the total read mass, even if their absolute abundance does not increase. This can produce statistically significant “up” calls that reflect relative reallocation rather than positive selection.

### 3.1.3 Main findings on the detected genes under selection

#### Analysis of the subset of 10 estimated essential genes

We present in Table 3 10 sgRNAs, which have smallest ratio 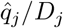 due to the bottleneck-neutrality test, and their corresponding genes together with the role of these genes. These are illustrated by the 10 red dots selection on the left-side of Figure 2.

The tabulated loci fall into three categories. Likely or near-essential determinants include *rplK-rplA, folA, rpsR-ssbA, nrdH-nrdE-nrdF*, and *murF*, which govern core translation, folate metabolism, DNA maintenance, nucleotide synthesis, and peptidoglycan biosynthesis. Conversely, *ylmG* is important for morphology and division but non-essential, as viable mutants persist despite division defects, while *glgA-glgD* encode accessory pathways for glycogen biosynthesis and carbon storage. The remaining loci (*pdxK, truA, pdxU2, tRNA-Tyr-2*, and *SPV 2330*) remain of uncertain essentiality. Since several sgRNAs target multi-gene operons, these assignments represent locus-level classifications rather than definitive single-gene calls.

On the genes most enriched (harmful to the pathogen)

The most significant enrichment signal is for sgRNA0223 (*p*_FDR_ = 2.51×10^*−*6^, lying on the horizontal dash-line in Figure 2), targeting *mtaR*, a *LysR*-family transcriptional regulator of streptococcal methionine metabolism and transport. Although methionine acquisition pathways influence *S. pneumoniae* growth and virulence Kovaleva and Gelfand (2007); Basavanna et al. (2013), *mtaR* enrichment in the lung may reflect niche-specific dispensability, as this regulon impacts nasopharyngeal persistence more heavily than pulmonary infection. Alternatively, given widespread library depletion, this signal could be a compositional artifact arising from the severe infection bottleneck skewing sgRNA representation. Thus, while a biological mechanism remains plausible, this result warrants caution absent individual mutant validation.

#### Link with Liu et al. (2021) findings and additional new insights

Remarkably, and reassuringly, all 265 of these sgRNAs that show down-behavior are also included in the list of 343 essential sgRNAs identified in the *in-vivo* mouse experiment, (Liu et al., 2021, Table S5). This implies that our method is **completely in line** with the method used in Liu et al. (2021), but it has the advantage that it narrows down even further the signal of selection to a smaller subset of sgRNAs.

#### Subsets of essential genes via statistical significance

Another point of view in selecting subsets of key essential genes, is by considering the *p*-values (Figure 2). Especially in our tests, many cases yield extremely small *p*-values ^5^, and one contributing reason is that the recipient sample has zero observed counts for one sgRNA. For an illustration of these non-neutral sgRNAs, we provide Table 4, describing notable sgRNAs strongly rejected by the test in either enriched or depleted directions (due to the *p*-value FDR-adjusted).

#### Genes that are non-neutral in-vivo but neutral in-vitro, and comparison with *purA*

To systematically identify niche-specific virulence factors that are exclusively required for host colonization rather than general metabolic maintenance, Liu et al. (2021) leveraged their dual-condition CRISPRi-seq platform to screen for differential fitness across environments. Among the candidate genes, the authors prioritized purA (encoding adenylsuccinate synthetase), which serves as a quintessential example of conditional essentiality. While purA remained entirely neutral during in vitro growth in rich C+Y medium, its repression led to a severe fitness defect within the murine respiratory tract. This gene emerged as the top hit in the screen, exhibiting both the largest magnitude of differential fitness (ΔFitness) in lung samples relative to the in vitro baseline and the highest statistical significance—measured by the−log_10_(*p*_adjusted_) of the interaction term (Liu et al., 2021, Figure 5). Mechanistic validation subsequently confirmed that this phenotype is driven by the stark scarcity of purine precursors within the host pulmonary environment, forcing the pathogen to rely strictly on de novo purine biosynthesis pathways.

We now compare the 265 depleted sgRNAs detected by our bottleneck-neutrality test, to the list of 309 essential genes in C+Y medium (Liu et al., 2021, Table S5), we have the 18 genes depleted in in-vivo but neutral in in-vitro ^6^. Among these sgRNAs, we compare the ratio between 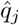 (estimated from Section 2.2 Step 4) and *D*_*j*_ (frequency of sgRNA *j* in the inoculum). The sgRNA0739 (target gene *srf-28*) has smaller ratio and *p*_FDR_ than these values of sgRNA0005 (target gene *purA*). In particular 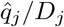 is 0.16023 for sgRNA0739 compared to 0.17951 for sgRNA0005. However, these values are quite close to each other, and this result also confirms the importance and neccessity of studying gene *purA* in Liu et al. (2021).

While *purA* operates as an established virulence factor, our analysis highlights *srf-28* as a newly suggested target with a distinct mechanistic consequence upon repression Liu et al. (2021); Slager et al. (2018); Weaver (2020). Specifically, *purA* encodes an adenylosuccinate synthetase critical for *de novo* AMP biosynthesis from IMP. Its repression does not cause immediate lethality in rich media but induces a conditional metabolic defect; silencing *purA* restricts nucleotide synthesis, growth, and replication within purine-poor *in vivo* niches, such as the murine lung. Conversely, *srf-28* is annotated as a putative Fst-family type I antitoxin sRNA within a toxin-antitoxin locus. Although its specific contribution to pneumococcal virulence requires experimental validation, repressing *srf-28* is predicted to be more acutely damaging than *purA* silencing. According to the Fst-family model, antitoxin loss derepresses the translation of the cognate toxin mRNA, producing a small, membrane-active peptide that immediately impairs cell-envelope function, cellular division, and growth, though direct evidence for this mechanism in pneumococcus remains outstanding.

### 4 Conclusion and discussion

The rapid expansion of high-throughput functional genomics—utilizing platforms such as genome-wide dual CRISPRi-seq to map epistatic networks Dénéréaz et al. (2025), cross-strain screens to evaluate evolutionary essentiality Rousset et al. (2020), or chemical genetics to profile drug-induced stress signatures Li et al. (2022); Sewgoolam et al. (2025), has generated exceptionally complex allele and sgRNA frequency datasets. Crucially, as these screens increasingly transition *in vivo* to study host colonization dynamics Maire et al. (2026), libraries routinely navigate strict experimental bottlenecks. Because these physical constrictions introduce massive stochastic variance (genetic drift), traditional differential abundance frameworks (e.g., DESeq2 or MAGeCK, see Supplementary material S3.1) risk compounding false discoveries. This operational landscape underscores the critical necessity for robust neutrality tests that can explicitly decouple bottleneck-driven drift from genuine selective pressure across high-throughput diversity data.

In this study, we propose a novel bottleneck-neutrality test that directly integrates transmission bottleneck size estimation into the statistical detection of selective pressure from biological diversity data. These data can consist namely of variant frequencies across two different points in time, or empirical conditions underlying a change in population size. By employing a Dirichlet-Multinomial sampling framework, building upon previous approaches (Sobel Leonard et al., 2017), our approach accounts for both the overdispersion inherent in sequencing data and the compositional constraint that variant frequencies must sum to unity.

Our application of the BN test to *in vivo Streptococcus pneumoniae* CRISPRi-seq data successfully isolated genotypic selection from stochastic loss. The framework not only replicated established essentiality findings, such as the *in vivo* fitness defect associated with *purA* repression and the core essentiality of *murF*, but also identified novel virulence candidates like the *srf-28* antitoxin. Furthermore, benchmarking against established differential abundance methods (for MAGeCK Li et al. (2014) and DESeq2 Love et al. (2014), see S3.2) highlighted a critical advantage of our approach: by explicitly anchoring the null hypothesis of neutrality to the physical population size parameter *N*_*b*_, the BN test correctly classifies stochastic frequency shifts caused by narrow founder populations as genetic drift. This mechanistic grounding prevents the false-positive selection calls prevalent in regression-based models that evaluate relative abundance without physical bottleneck constraints.

#### Comparison of BN test with other methods

We compare our proposed bottleneck-neutrality (BN) test with two widely adopted frameworks, MAGeCK and DESeq2 (see Table S2 for a brief overview of each methodology). All three methods analyze high-throughput sequencing read counts and must account for over-dispersion, where the variance of biological count data exceeds its mean. They therefore avoid simple Poisson models and use probability models that capture extra biological and sampling variability. The differences between BN test with MAGeCK/DESeq2 are presented in the following Table 5. The choice between DESeq2, MAGeCK, and the BN test depends on the data type and experimental dynamics. DESeq2 is appropriate for standard transcriptomics (e.g., RNA-seq), where its Negative Binomial model and empirical Bayes shrinkage effectively resolve overdispersion in static mRNA expression. However, it is unsuitable for pooled genetic screens, particularly bottlenecked *in vivo* models, because its lack of physical transmission/population size parameters causes it to misinterpret stochastic variant loss as directional selection. For pooled screens, tool selection depends on the magnitude of genetic drift. MAGeCK is optimal for *in vitro* and non-bottlenecked *in vivo* models, where large founding populations render drift negligible. Conversely, for *in vivo* models with narrow transmission bottlenecks, the BN test should be favoured (e.g. see Section S3.2). By incorporating the estimated bottleneck size directly into a Beta-Binomial distribution, the BN test mathematically bounds stochastic variance, successfully separating niche-specific adaptation from founder-effect drift.

**Table 5:**
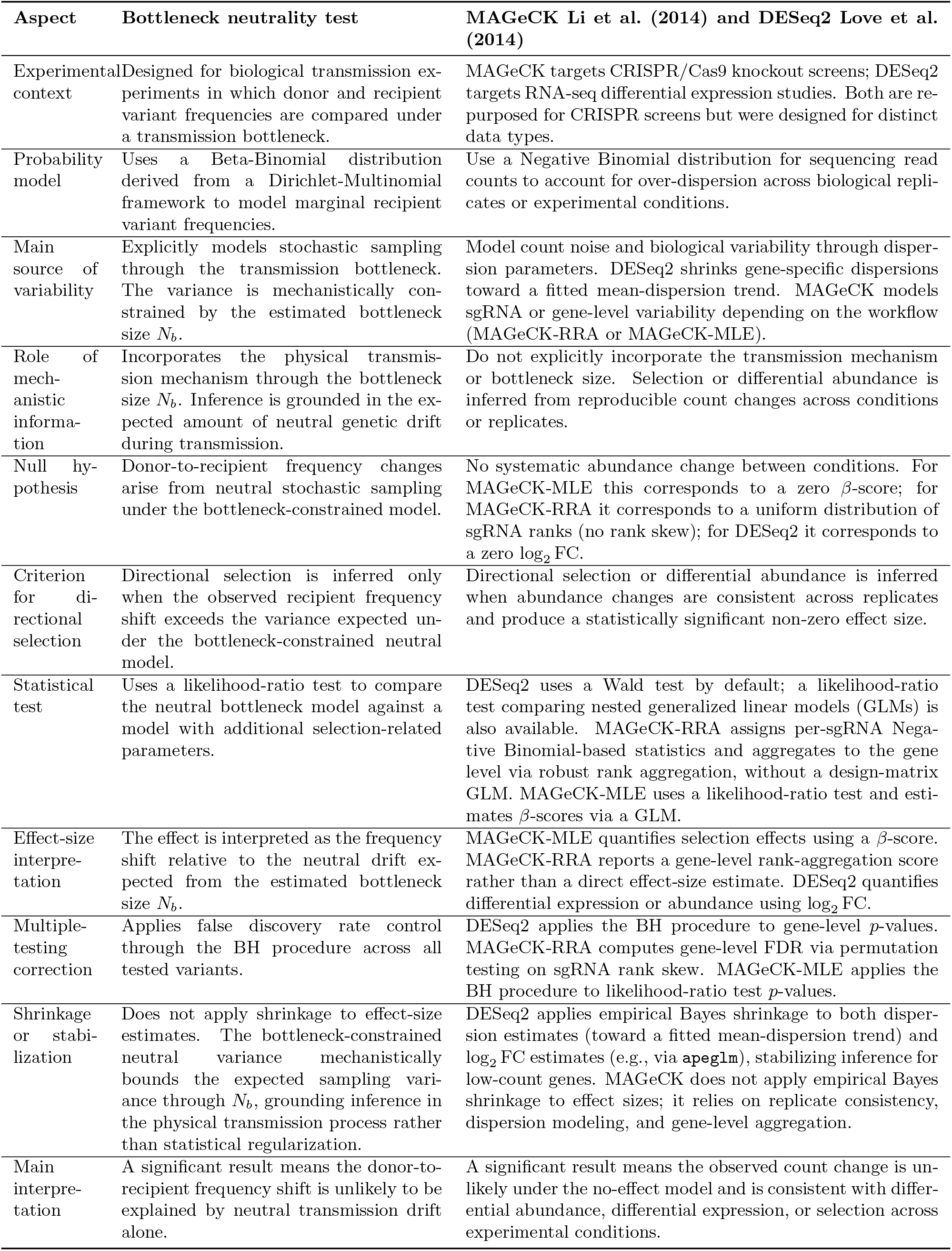
Comparison between the bottleneck neutrality test and count-based differential abundance methods (MAGeCK and DESeq2).

#### Limitations of BN test and directions for extension

The BN test is subject to two interrelated mathematical limitations arising from data compositionality and the majority-neutrality assumption. First, because sequencing reads represent relative frequencies that sum to unity, variant abundances are statistically dependent. A large positive selective sweep on a subset of variants mathematically forces the dilution of remaining frequencies, potentially generating false-positive signatures of negative selection among neutral alleles. Secondly, the Dirichlet-Multinomial fitting assumes that the majority of the library is neutral. When widespread selection violates this assumption, it inflates the observed variance across variants. The model mis-attributes this excess variance to stochastic drift rather than selection, causing *N*_*b*_ to be underestimated. A deflated *N*_*b*_ widens the neutral noise margin in the likelihood-ratio test, reducing statistical power and increasing false negatives as genuine selection signals fall within the inflated drift boundaries.

Together, these limitations make the BN test susceptible to compositional false positives under strong directional selection and to false negatives under widespread selection.

Despite these limitations, incorporating physical bottleneck sizes into likelihood-ratio testing provides a novel mechanistically grounded tool for microbial genomics. This integrated model improves the precision and interpretability of fitness screens, establishing a robust framework to characterize pathogen transmission, infection dynamics, and evolutionary selection across diverse systems.

Several extensions could address the identified limitations and broaden the framework’s applicability. The majority neutrality assumption in the Dirichlet-Multinomial fitting could be relaxed by estimating *N*_*b*_ on a pre-filtered neutral subset of variants, analogous to using non-targeting controls for dispersion estimation in MAGeCK, reducing the influence of widespread selection on bottleneck calibration. Compositional false positives could be mitigated by rescaling recipient frequencies against a known-neutral reference before applying the LRNT, decomposing reallocation artefacts from genuine depletion. Inter-host heterogeneity in *N*_*b*_ could be captured through a hierarchical extension that places a prior over *N*_*b*_ across hosts and marginalizes over estimation uncertainty, yielding better-calibrated *p*-values when host numbers are small or bottleneck sizes vary substantially. Furthermore, accounting explicitly for uncertainty in estimates of population bottleneck size, and how that uncertainty affects the selection test, remains an important extension for the future. Overall many directions for methodological improvement exist, making the Bottleneck-Neutrality framework a flexible and generalizable tool for better quantification of selection processes across many biological systems.

## Code availability

GitHub: https://github.com/lthminhthao/Integrating-Bottleneck-Size-Estimation-into-Selection-Tests

## Acknowledgements

We thank Jan-Willem Vening, Jean-Claude Sirard and Absalom Janssen for helpful discussions.

## Funding

This work was funded by the European Union (Grant agreement 101080528 NOSEVAC project).

## Supplementary Information

### S1 Bottleneck size estimation

#### S1.1 The binomial model

The binomial model infers bottleneck size by treating transmission as a probabilistic sampling process where each transmitted individual represents an independent draw from the donor population, meaning the number of inherited variant copies follows a binomial distribution. While Sobel Leonard et al. (2017) is a seminal genomic application of this framework to estimate transmission bottlenecks, earlier foundational work by (Emmett et al., 2015) and (Poon et al., 2016) similarly utilized binomial-type variant frequency models in pathogen transmission contexts. The number of transmitted individuals carrying the variant *k*_*i*_ follows *k*_*i*_ ~ Binomial (*N*_*b*_, *D*_*i*_). After transmission, the recipient population is assumed to expand, and the observed recipient frequency reflects *R*_*i*_ = *k*_*i*_*/N*_*b*_, possibly with additional sampling noise from sequencing. The likelihood of bottleneck can be written as:

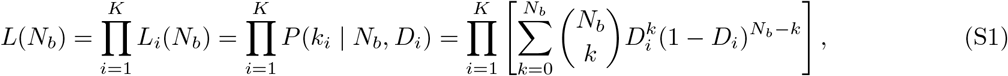

where *P* (*k*_*i*_ | *N*_*b*_, *D*_*i*_) denotes the binomial distribution evaluated at *k*_*i*_ and parameterized with *N*_*b*_ number of trials and a success probability of *D*_*i*_. The bottleneck size *N*_*b*_ can be estimated using maximum likelihood by choosing *N*_*b*_ that maximizes *L*(*N*_*b*_), or Bayesian inference by placing a prior distribution on *N*_*b*_. To use this binomial model, we need these additional assumptions

- Independent sampling of individuals from the donor population, i.e. independence among variant sites primarily to ensure the mathematical tractability of the global likelihood function.
- Post-transmission genetic drift is ignored or simplified.

While analytically tractable and effective for narrow bottlenecks, the binomial model neglects donor frequency uncertainty and sequencing errors, introducing bias under wide bottlenecks or substantial post-bottleneck drift. Consequently, it is often extended via beta-binomial distributions to capture overdispersion in applications like viral transmission, experimental evolution, and founder events.

#### S1.2 Proof of the bottleneck size computed used beta-binomial model

*Proof*. We explain (2.1) and (2.2) in Section 2.1. To obtain the overall likelihood of population bottleneck size *N*_*b*_, we consider all possible scenarios of how many virions out of the total *N*_*b*_ virions transferred carried the variant allele.

The probability that the founding population of *N*_*b*_ virions carries *k* variant alleles is given by the binomial distribution

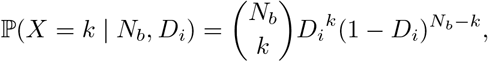

where the number of trials is *N*_*b*_ and the success probability is *D*_*i*_, the frequency of variant *i* in the donor.

For a stochastic birth–death process with constant birth rate *λ* and death rate *µ*, the probability mass function for the viral population size originating from a single virion that successfully establishes infection follows the geometric distribution

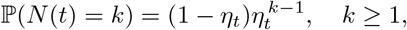

where *t* is the time of sampling, *N* (*t*) is the population size at time *t*, and 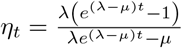 due to (Kendall, 1948, (15) & (16)).

Indeed, according to Kendall (1948), the population sizes stemming from each of the *N*_*b*_ founding virions—each having successfully established infection—are thus geometrically distributed random variables. As these population sizes are likely to be very large at the time of sampling, we can approximate them, which follows geometric distribution, as being exponentially distributed random variables.

Under this approximation, the distribution of the fractions of the population descended from each of the founding virions follows a Dirichlet distribution: Dirichlet(1, 1, …, 1), with *N*_*b*_ components, one for each ancestor. Indeed, we have this well-known result.

##### Lemma 1.

*Suppose X* = (*X*_1_, …, *X*_*n*_) *are independent, identically distributed random variables with X*_*i*_ ~ Exp(*λ*), *that is, f*_*Xi*_ (*x*) = *λe*^*−λx*^**1**_[0,*∞*)_(*x*). *Setting Y* = (*Y*_1_, …, *Y*_*n*_), *where* 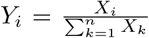 *then Y follows a Dirichlet distribution Y* ~ Dir(1, 1, …, 1).

A subset *k* of these founder virions carries the variant allele, while the remaining (*N*_*b*_−*k*) founders carry the reference allele. We recall another well-known result in (Johnson et al., 1972, p. 489).

##### Lemma 2.

*Let X* = (*X*_1_, …, *X*_*n*_) *be a random vector following a Dirichlet distribution X* ~ Dir(*α*_1_, …, *α*_*n*_) *with α*_*i*_ *>* 0. *Let us partition the components of X into two disjoint sets and define a new random variable Y as the sum of the first k components of X (where* 1 ≤ *k < n):* 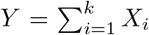 *Consequently, the sum of the remaining components is* 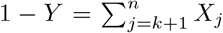 *Then the marginal distribution of the sum Y follows a Beta distribution* 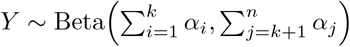

Collapsing the Dirichlet distribution yields that the fraction of the population carrying the variant allele follows a Beta distribution Beta(*k, N*_*b*_ − *k*). Thus, we conclude (2.1) and (2.2).

#### S1.3 Bottleneck size in Abel et al. (2015)

Abel et al. (2015) established a framework to estimate infection bottleneck sizes (*N*_*b*_), defined as the number of founding lineages surviving a population constriction. By introducing a diverse inoculum of genetically barcoded microbes and tracking frequency shifts post-expansion via high-throughput sequencing, the method captures bottleneck-induced genetic drift. Specifically, the framework applies population genetics estimators from Krimbas and Tsakas (1971) to relate the variance in barcode frequency changes directly to an effective population size, yielding the estimated bottleneck size.

Concretely in Abel et al. (2015), the size of bottleneck is calculated as

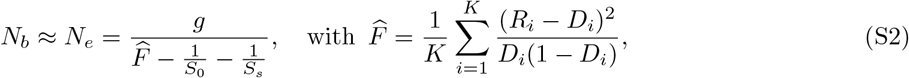

recalling *K* is the total number of distinct alleles, and denoting *g* the number of generations during competitive growth, and *S*_0_ and *S*_*s*_ the sample sizes used to determine the population composition at time 0 and at sampling, respectively. The approach also assumes the selective neutrality of barcodes.

To ensure accuracy, Abel et al. (2015) calibrated raw effective population size estimates against *in vitro* controls with known bottleneck sizes, defining *N*_*b*_ as the number of surviving founding lineages. Grounded in classical population genetics theory, this framework leverages high barcode diversity and deep sequencing to quantify bottleneck-induced drift with high resolution across a wide dynamic range.

*Proof*. For each variant *i*, recalling the observed frequencies (from sequencing) be *D*_*i*_, *R*_*i*_ at time 0 and time *s*, respectively, then we denote that

- *p*_*i*,0_ and *p*_*i,s*_ are population frequencies of variant *i* after the drift through bottleneck, in the inoculum (time 0) and at the sampling time *s* (after *g* generations of within-host growth), respectively.
- *X*_*i*,0_ is the number of reads at time 0 that is assigned to allele *i*, then *X*_*i*,0_ ~ Bin(*S*_0_, *p*_*i*,0_) and *D*_*i*_ =*X*_*i*,0_*/S*_0_.
- Decompose the observed change as: *R*_*i*_ − *D*_*i*_ = (*p*_*i,s*_ − *p*_*i*,0_) + (*R*_*i*_ − *p*_*i,s*_) − (*D*_*i*_ − *p*_*i*,0_) = Δ*p* + *ε*_*s*_ − *ε*_0_.
- Under neutrality and independence of errors, we have that E[Δ*p*] = 0, E[*ε*_0_] = 0, E[*ε*_*s*_] = 0, so E[*R*_*i*_ −*D*_*i*_] = 0. Under independence of drift and sampling errors (and of the two sampling errors), (Δ*p, ε*_0_, *ε*_*s*_) independent then E[Δ*p ε*_*s*_] = E[Δ*p*] E[*ε*_*s*_] = 0, and similarly E[Δ*p ε*_0_] = 0, E[*ε*_*s*_ *ε*_0_] = 0. By direct computation: E(*R*_*i*_ − *D*_*i*_)^2^ ≈ Var(Δ*p*) + Var(*ε*_*s*_) + Var(*ε*_0_).

We now analyze each term.

1. Drift through the bottleneck (effective size *N*_*b*_, *g* generations). For a neutral Wright–Fisher–type process starting at *D*_*i*_, and drift over *g* generations with effective *N*_*b*_, the diffusion approximation gives

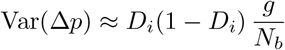

for moderate *g* and small *g/N*_*b*_. This is the “evolutionary” piece; variance grows linearly with *g/N*_*b*_. Indeed, for a neutral Wright–Fisher process, by Falconer (1981), one can derive

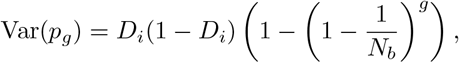

and when *g* ≪ *N*_b_, using the approximation 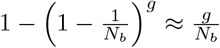allow us to recover exactly

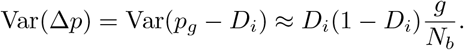

more generations or a smaller founding population both increase the variance in frequencies due to neutral drift.
2. Sampling at time 0 (sequencing depth *S*_0_): counts are binomial, so 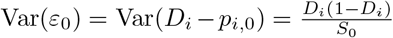 Indeed, for the binomial random variable *X*_0_, E[*X*_0_] = *S*_0_*D*_*i*_ and Var(*X*_0_) = *S*_0_*D*_*i*_(1 − *D*_*i*_). Then

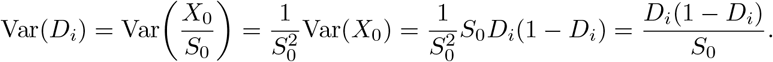

Since *X*_*i*,0_ ~ Bin(*S*_0_, *p*_*i*,0_), implying E[*X*_*i*,0_] = *S*_0_*p*_*i*,0_, then E[*ε*_0_] = E[*D*_*i*_] − *p*_*i*,0_ = E[*X*_*i*,0_*/S*_0_] − *p*_*i*,0_ = E[*X*_*i*,0_]*/S*_0_ − *p*_*i*,0_ = 0. Then Var(*ε*_0_) = Var(*D*_*i*_), which gives the stated formula.
3. Sampling at time *s*: similarly, using *p*_*g*_ ≈ *D*_*i*_ for small changes 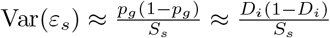 Hence, we have that E[(*R*_*i*_ − *D*_*i*_)^2^] ≈ *D*_*i*_(1 − *D*_*i*_) ^*g* 1^ + ^1^. We have the statistic *F*^ normalized by *D*_*i*_(1 − *D*_*i*_) and then the expectation:

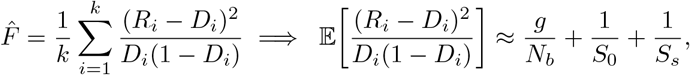

and averaging over *k* alleles does not change this expectation.

### S2 Application to data for bottleneck size estimation

#### S2.1 Summary of Liu et al. (2021): Exploration of Bacterial Bottlenecks

Liu et al. (2021) applied a genome-wide CRISPR interference sequencing (CRISPRi-seq) approach in *Streptococcus pneumoniae* D39V to simultaneously characterize infection bottlenecks and identify genes required for host survival and dissemination. The authors constructed a pooled, doxycycline-inducible library of 1,499 unique sgRNAs targeting specific operons. Following *in vitro* validation in C+Y medium, mice were challenged with the pool, and sgRNA abundances were quantified via Illumina sequencing. The underlying raw counts across the inoculum, uninduced, and induced groups for each animal at 24 and 48 hpi are detailed in Table S7 (corresponding to Figures 3-6).

#### S2.2 Replicating results from Liu et al. (2021) and application of the Dirichlet-multinomial method

Because sgRNA frequencies sum to unity, the site-independence assumption of the beta-binomial model is violated, we therefore employ the DM model described in Section 2.1 to estimate transmission bottleneck sizes and compare our results with the founding population sizes reported by Liu et al. (2021).

**Figure S1:**
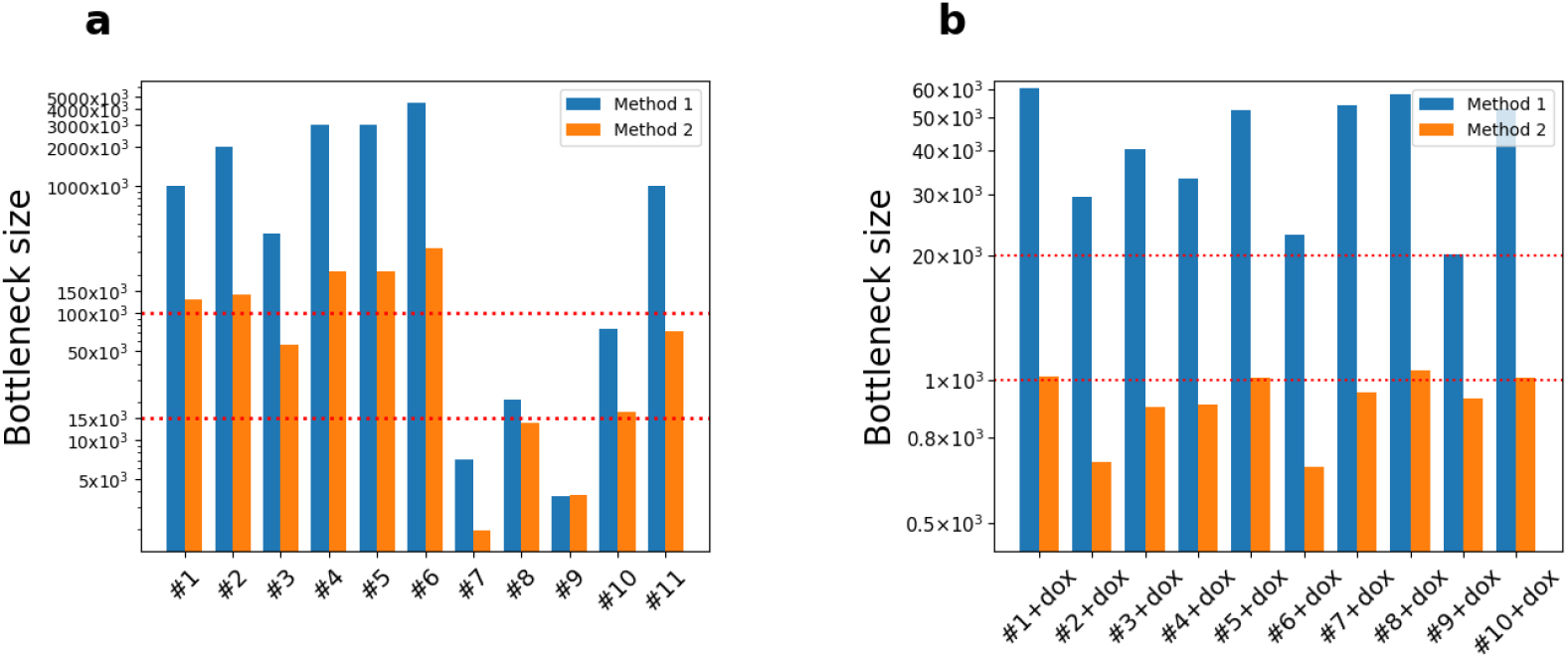
Comparison of foundation population and transmission bottlenecks under no-dox and with-dox conditions at 24 hpi. Paired bars compare estimated population bottleneck sizes (blue; Section S1.3) and transmission bottleneck sizes (orange; Section 2.1) for individual mice under (a) uninduced (no-dox) and (b) induced (+dox) conditions. The panels utilize a piecewise, non-linear *y*-axis to display values spanning multiple orders of magnitude. Across both conditions, transmission bottlenecks are consistently lower than population bottlenecks.

The order-of-magnitude discrepancy between the two estimates reflects fundamental differences in their biological and mathematical formulations. The framework in Section S1.3 treats *N*_*b*_ as an effective population size (*N*_*e*_) integrated over the growth period Abel et al. (2015). Because this estimate scales linearly with the number of generations—calculated based on an *S. pneumoniae* doubling time of 108 minutes—the extensive bacterial expansion by 24 hpi inflates *N*_*b*_. This value therefore reflects the attenuated genetic drift of an expanded population rather than the physical count of the initial founding seed Liu et al. (2021). Conversely, the DM framework captures the transmission bottleneck, yielding substantially lower values of approximately 10^5^ at 24 hpi and 80 at 48 hpi (excepting mouse 12, with an estimated *N*_*b*_ of 200 in the lung and 40 in blood).

Similarly, we have the similar observation in the comparison of foundation population and transmission bottleneck sizes at 48 hpi in lung and blood as follows in Figure S2.

Mechanistically, the framework in Liu et al. (2021) captures a post-expansion effective population size (*N*_*e*_) rather than the number of initial founding bacteria. Conversely, transmission bottleneck estimation determines the number of independent founders required to reproduce the frequency dispersion observed in the recipient data. Because the statistical signature of a finite founder population remains encoded in the variance structure of downstream frequencies, this maximum likelihood estimation (MLE) procedure is insulated from post-bottleneck growth, reflecting the founding population scale more directly than approaches that scale explicitly with the number of generations *g*.

To evaluate whether the population-to-transmission bottleneck ratio 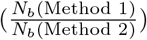 corresponds to this growth factor, we analyzed these ratios across all condition, tissue, and timepoint groups. A one-sample, two-sided Student’s *t*-test was applied to assess significant deviations from the *a priori* reference values of *g* (13.333 at 24 hpi; 26.666 at 48 hpi).

From the Table S1, except the case of 48 hpi Lung of mice with Doxycycline, we reject the hypothesis that the observed bottleneck ratio is equal to the number of generation *g*. To visualize these statistical values, we have the following summary figure.

**Figure S2:**
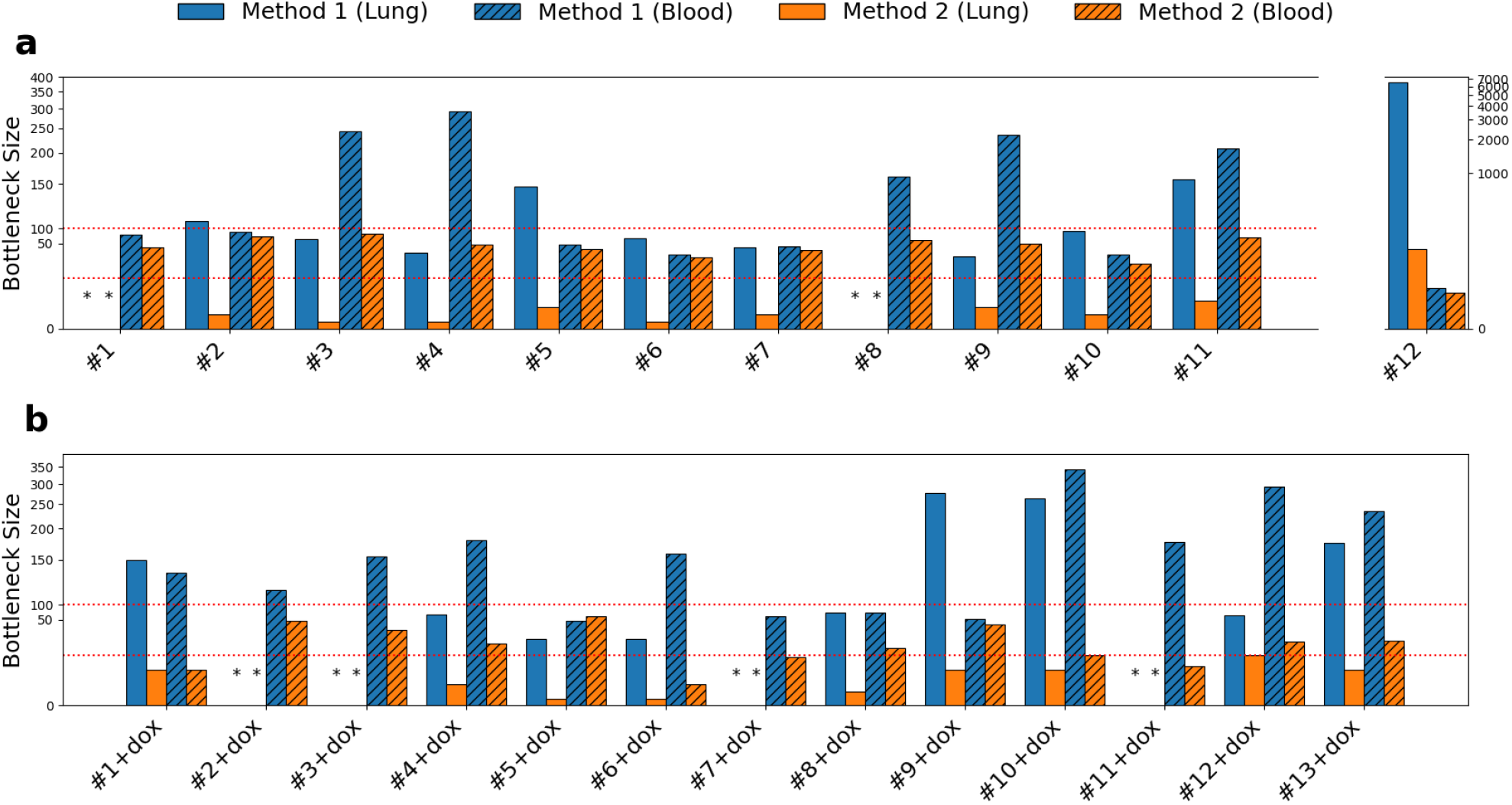
Comparison of foundation population and transmission bottlenecks under no-dox and **with-dox** conditions at 48 hpi. Bar plots comparing bottleneck size estimates at 48 hours post-infection (hpi) in lung and blood for individual mice, contrasting population bottleneck size *N*_*b*_ (blue; Section S1.3) with transmission bottleneck size *N*_*b*_ (orange; Section 2.1). Lung bars feature a solid fill and blood bars are hatched. Panels display mice **(a)** without doxycycline, with mouse #12 shown in a separate side panel to accommodate its larger range, and **(b)** with doxycycline, where black asterisks denote lung samples without bacterial recovery. Across both panels, transmission bottlenecks are lower than population bottlenecks, though less pronouncedly than in Figure S1.

**Figure S3:**
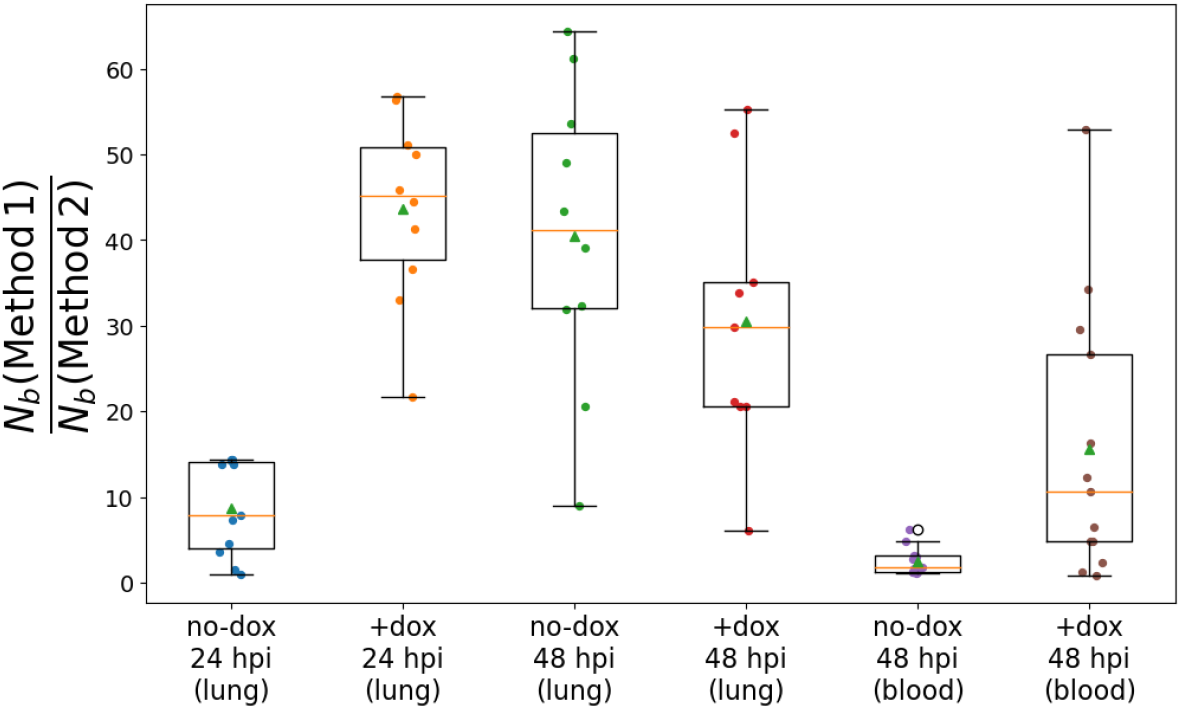
Summary of population-to-transmission bottleneck ratios across timepoints, tissues, and **induction** conditions. Box-and-whisker plots showing individual *N*_*b*_(Method1)*/N*_*b*_(Method2) ratios grouped by condition, tissue, and timepoint. Dots represent individual mice, triangles denote group means, box centers indicate medians, and whiskers extend to 1.5 *×* IQR. Ratios are consistently higher in the lung than in blood—peaking under +dox at 24 hpi—whereas blood at 48 hpi displays the lowest ratios, indicating closer agreement between the two bottleneck estimators in circulation than in lung tissue.

### S3 MAGeCK and DESeq2

#### S3.1 Summary of MAGeCK and DESeq2

This supplementary section provides an overview of the MAGeCK and DESeq2 methodologies, which are presented in the following Table S2.

**Table S1:** One-sample two-sided *t*-test results comparing *N*_*b*_(Method 1)*/N*_*b*_(Method 2) ratios to the reference values (13.333 at 24 hpi; 26.666 at 48 hpi).

| Timepoint | Tissue | Group | $n$ | Mean | $t$-stat | $p$ | 95% CI |
| --- | --- | --- | --- | --- | --- | --- | --- |
| 24 hpi | Lung | noDox | 11 | 8.76 | -2.75 | $2.05 \times 10^{-2}$ | [5.06, 12.47] |
| 24 hpi | Lung | Dox | 10 | 43.69 | 8.75 | $1.08 \times 10^{-5}$ | [35.84, 51.54] |
| 48 hpi | Lung | noDox | 10 | 40.45 | 2.48 | $3.50 \times 10^{-2}$ | [27.88, 53.02] |
| 48 hpi | Lung | Dox | 9 | 30.54 | 0.73 | $4.84 \times 10^{-1}$ | [18.35, 42.73] |
| 48 hpi | Blood | noDox | 12 | 2.49 | -51.57 | $1.79 \times 10^{-14}$ | [1.46, 3.52] |
| 48 hpi | Blood | Dox | 13 | 15.62 | -2.52 | $2.70 \times 10^{-2}$ | [6.07, 25.18] |

#### S3.2 Technical walk-through (benchmarking rationale)

To demonstrate why omitting the *N*_*b*_ parameter impairs MAGeCK and DESeq2 when evaluating in-vivo transmission data, we consider the following example.

##### Data configuration and MAGeCK / DESeq2 processing

The baseline frequencies of variant *j* in the inoculum (donor) and in the recipient are *D*_*j*_ = 0.05 and *R*_*j*_ = 0.15, respectively. The total sequencing depth for the recipient host is *n*_*m*_ = 10^5^.

Both MAGeCK and DESeq2 evaluate this change via the log2FC: log_2_(0.15*/*0.05) ≈1.58. Given the high sequencing depth (corresponding to large absolute read counts of 5,000 and 15,000) and assuming low technical dispersion between replicates, these Negative Binomial models will return a highly significant *p*-value. They flag variant *j* as exhibiting “differential abundance” because the algorithms are blind to the physical bottleneck constraints of the transmission event.

#### BN test technical inference (leveraging *N*_*b*_)

Evaluating the exact same frequency shift (0.05→0.15) under the Beta-Binomial framework, and assuming negligible technical dispersion, the theoretical variance of a variant’s recipient frequency relative to its baseline donor frequency is approximated by 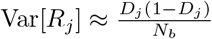 We consider the following two cases:

##### 1. Narrow Bottleneck

*N*_*b*_ = 20: 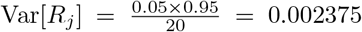, yielding a standard deviation of *Σ* ≈ 0.0487. An observed 10% frequency increase represents a 2.05*Σ* deviation. Because small founder populations generate substantial stochastic uncertainty (founder effect), the algorithm classifies this shift as random sampling noise, thereby retaining *H*_0_ and preventing a false-positive selection call.

##### 2. Wide Bottleneck *N*_*b*_ = 5000

The variance decreases to 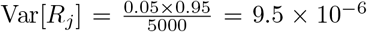, yielding a standard deviation of *Σ* ≈0.00308. The identical 10% shift constitutes a 32.4*Σ* deviation, which is statistically implausible under neutral dynamics since random drift is minimized in a large population. The LRNT thus rejects *H*_0_, confirming deterministic positive selection.

**Table S2:**
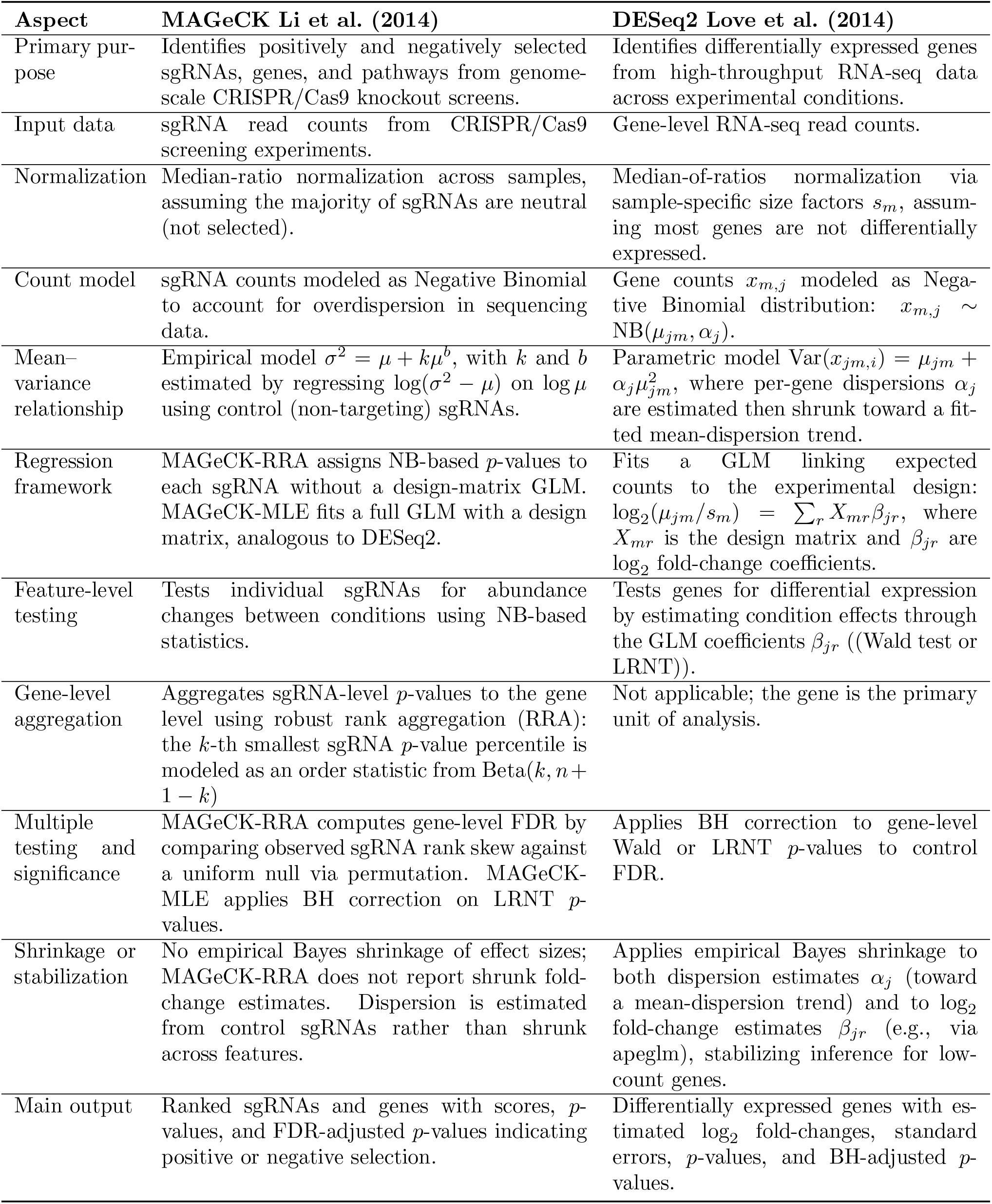
Summary and comparison between MAGeCK and DESeq2.

## Footnotes

1 At 24 hpi, *N*_*b*_ spanned [2 ×10^4^, 6×10^4^] for the doxycycline-treated group, whereas the control group exhibited larger bottlenecks, with seven mice in the range [10^6^, 5 X10^6^] and two in [2 X10^4^, 8 X10^4^]. By 48 hpi, bottlenecks constricted severely across both groups into an overall range of [25, 300]—falling below 300 in blood and 200 in lung for all animals except control mouse 12—with no significant difference between conditions.

2 **Results:** Screening identified 339 sgRNAs targeting essential genes or operons *in vitro* (C+Y medium) and 343 *in vivo* in the pneumonia model (lung, 24 hpi).

3 **Results:** Comparative screening identified 46 sgRNAs with significant differential fitness (|ΔFitness| *>* 1, *p*_adj_ *<* 0.05), included 31 targets more essential *in vivo* (e.g., *purA* and the *cps* capsule locus) and 15 more essential *in vitro* (e.g., *metK*).

4 sgRNA0396 (down, target: nrdH, nrdE, nrdF), sgRNA1347 (up, target: hexA), sgRNA0501 (down, SPV 1350, target: unknown), sgRNA0492 (down, target: tuf), sgRNA0081 (down, target: rpsE, rplX, secY, rpsN, rplN, rplR, rpmD, rplF, rpsQ, rplO, rpmC, rplE, rpsH), sgRNA0406 (up, SPV 1072, target: unknown), sgRNA0575 (up, SPV 2382, target: unknown), sgRNA0525 (down, target: dpr), sgnRNA1457 (up, SPV 2279, target: unknown)

5 For example, in mouse 1 with dox, we have the following sgRNAs that cannot be read at 24 hpi: sgRNA765, sgRNA860, sgRNA501, sgRNA676, sgRNA492, sgRNA1342, sgRNA1340, sgRNA283, sgRNA302, sgRNA549.

6 sgRNA0003, sgRNA0005, sgRNA0206, sgRNA0289, sgRNA0370, sgRNA0374, sgRNA0464, sgRNA0475, sgRNA0525, sgRNA0532, sgRNA0606, sgRNA0706, sgRNA0739, sgRNA0856, sgRNA1094, sgRNA1239, sgRNA1308, sgRNA1341

## Notes

https://www.cell.com/cell-host-microbe/fulltext/S1931-3128(20)30559-X?_returnURL=https%3A%2F%2Flinkinghub.elsevier.com%2Fretrieve%2Fpii%2FS193131282030559X%3Fshowall%3Dtrue

### Competing Interest Statement

The authors have declared no competing interest.

